# Prevalence and Associated Risk Factors of Bovine Trypanosomosis in Diga District, Western Ethiopia

**DOI:** 10.1101/2024.02.11.579830

**Authors:** Abebe Tesfaye Gessese, Gemechis Yadesa

## Abstract

Trypanosomosis is a protozoal disease caused by different species of unicellular parasites found in the blood and other tissues of vertebrates including livestock, wildlife, and people. It is the most serious animal production problem in sub-Saharan Africa including Ethiopia. The present study was conducted to assess the prevalence of bovine trypanosomosis and prevailing species of Tyrpanosomes in Diga district and to identify host related risk factors for trypanosomosis prevalence in Western Ethiopia. Blood samples were collected from 384 randomly selected indigenous cattle (*Bos indicus*) and analyzed using parasitological and hematological techniques. Out of the 384 animals examined 22 (5.73%) were found trypanosomosis positive. Of which 11 (2.87%) were infected with *T. vivax*, 9 (2.34%) with *T. congolense* and 2 (0.52%) with mixed infection of *T. vivax* and *T. congolense*. This difference in trypanosomes species prevalence was not statistically significant (P>0.05). The association of prevalence with the various risk factors including kebele, age, sex, body condition and coat color were measured. There was a significant variation in the prevalence of trypanosomosis (P<0.05) among cattles with different body conditions, poor body conditioned animals with the highest prevalence followed by medium and good body condition animals, respectively. There was no any statistically significant difference in other variables considered. The overall Packed Cell Volume (PCV) mean PCV value of 25.67 was recorded during the study period. The mean PCV value of infected animals was lower (20.18) than that of non-infected animals (26.0). The difference was statistically significant (P<0.05). The result of the present study indicated the prevailing occurrence of trypanosomosis in the study area necessitating integrated control measures.

## INTRODUCTION

Trypanosomosis is the most serious veterinary and animal production problem in sub-Saharan Africa and prevents the keeping of ruminants and equines over 10 millions of square kilometers of potentially productive land. Hence, the road map and contribution to the Pan African Tsetse and Trypanosomosis Eradication Campaign agenda [1]. It is a protozoal disease caused by different species of unicellular parasites found in the blood and other tissues of vertebrates including livestock, wildlife, and people [2]. Tsetse flies inhabit wide range of habitats covering over 10 million Km_2_ representing 37% of the African continent and affecting 38 countries [3] including Ethiopia. They are restricted to various geographical areas according to habitat, the three main groups, named after the commonest species in each group, being *fusca, palpalis*and *morsitans*, found respectively in forest, riverine and savannah areas. The last two groups, because of their presence in the major livestock rearing areas, are the most from a veterinary standpoint [4, 5]. In Ethiopia, they are confined southern and western regions between longitude 330 and 380 E and latitude 50 and 120 N which amounts to about 200,000 km_2_. Tsetse infested areas lied in lowlands and in river valley of Abay (Blue Nile), Baro, Akobo, Didessa, Ghibe and Omo. Out of the nine regions of Ethiopia five (Amhara, Benishangul-gumuz, Gambella, Oromia and SNNPR) are infested with more than one species of tsetse flies [6, 7].

Tsetse transmitted animal trypanosomosis is one of the most significant and costly disease in the country where tsetse flies are highly distributed, hindering the effort being made for food self-sufficient. In Ethiopia about 200,000km^2^ of the land is infested with tsetse flies and preclude farmers from rearing livestock. Disease could affect development through its historical effect on shaping institutions and/or through contemporaneous impacts on health [8]. Another indirect effect on the economic development of the country is the costs of drug to treat the disease and control of the tsetse flies. The added risk of human infections due to sleeping sickness, the most fatal trypanosome disease transmitted by tsetse fly has also greatly affected social, economic, and agricultural of the rural communities [5, 9].

In the tsetse-infested areas of Africa, trypanosomiasis is well recognized, and diagnosis is often based on history of chronic wasting condition of cattle in contact with tsetse flies. Trypanosome can be confirmed parasitologically by demonstrating parasites in blood of infected animals and various techniques are available. In practice, many field programs of monitoring cattle for infection is based on routine screening of stained thick and thin blood films, thick films are examined to detect infected animals and thin films determine the species of infecting trypanosomes [10].

Therefore, the objectives of the present study were to assess the prevalence of bovine trypanosomosis and prevailing species of Tyrpanosomes and identify host related risk factors for trypanosomosis prevalence in Diga district.

## 3. MATERIALS AND METHODS

### 3.1. Study Area

The study was conducted from November 2019 to March 2020 in Diga district of East Wollega, Oromia regional state, Ethiopia. Diga is located 345 km distance from Addis Ababa and 12 km from the capital city of East Wollega zone, Nekemte, at altitude of 2250 masl. It receives annual average rain fall of approximately 1250mm. The annual temperature varies from 14°C – 32°C with average temperature 22.6°C [11]. There are two rainy seasons namely the summer and spring. The rain shower falling in the spring season was medium which was very important for growth of plants as whole. The main rain season for the study area was summer whereby sufficient rain and moisture was available for plant growth [12]. The district is characterized by crop livestock mixed farming system. Teff, Wheat, barley, maize, sorghum, peas, beans, chickpea, linseeds, Nug and rape seed are the major annual crops grown in the area. The estimated animal population of the area was 67,060 cattle among cattle population 15 pure Borena breed, 144 cross breed (holystein feresian with zebu breed), 11,893 sheep, 6,426 goats, 3066 donkey, 147 horses and 48 mules [11].

### 3.2. Study Population

The study animals were zebu cattle (Bos indicus) kept under extensive traditional husbandry condition grazing communally owned pastureland throughout the year. Total of 384 cattle were sampled at their communal grazing area. The body condition of each of the study cattle was scored as good, medium, and poor. Simultaneously, their age was determined based on dentition [13].

### 3.3. Study Design

A cross sectional study was conducted from November 2019 to March 2020 in order to assess the prevalence of bovine trypanosomosis and identify host related risk factors for trypanosomosis prevalence in Diga district.

### 3.4. Sample size determination and samplind strategy

The number of animals required for the study was determined using the formula given by

[14] for simple random sampling.

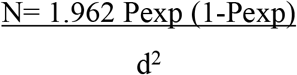

Where N = required sample size

P exp= expected prevalence

d = desired absolute precision (usually 0.05)

The size of the sample is determined using 95% level of confidence, 50% expected prevalence, since there was no similar study in the study area, and 0.05-desired absolute precision. Therefore, 384 cattle’s were needed for the study.

Animals were selected from the study population using simple random sampling technique and purposive sampling was the method followed to select the three study peasant associations (Kebeles).

### 3.5. Sample collection and procedures

To determine the Packed Cell Volume (PCV), Blood samples were obtained by puncturing the marginal ear vein with a lancet and collected directly into heparinized capillary tubes. The tubes were then sealed at one end with crystal seal. PCV was measured in a micro haematocrit centrifuge. The capillary tubes were placed in microhaematocrit centrifuge with sealed end outer most. Then the tube was loaded symmetrically to ensure good balance. After screwing the rotary cover and closing the centrifuge lid, the specimens were allowed to centrifuge at 12,000 rpm for 5 minutes. After centrifugation, the capillary tubes were placed in a haematocrit reader. The length of the packed red blood cells column is expressed as a percentage of the total volume of blood. Animals with PCV less than 24% were considered to be anaemic.

#### 3.5.1 Buffy coat technique

Heparinised microhaematocritcapillary tubes, containing blood samples were centrifuged for 5 minutes at 12,000 rpm. After the centrifugation, trypanosomes were usually found in or just above the buffy coat layer. The capillary tube was cut using a diamond tipped pen 1 mm below the buffy coat to include the upper most layers of the red blood cells and 3 mm above to include the plasma. The content of the capillary tube was expressed onto a glass slide and covered with cover slip. The slide was examined under x40 and x10 eye piece for movement of parasite. Trypanosome species were identified according to their morphological descriptions as well as movement in wet film.

### 3.6. Data Analysis

All the collected raw data were entered into a Microsoft excel spread sheets program and then was transferred to Stata/MP version 16 for analysis. The prevalence of trypanosome infection was calculated as the number of positive animals examined by buffy coat method divided by the total number of animals examined at the particular time. Pearson’s chi-square was used to evaluate the association of different variables with the prevalence of trypanosome infection. In all the statistical analysis executed, a confidence level of 95% is used and P-value of less than 0.05 (at 5% level of significance) was considered as statistically significant.

## 4. RESULTS

### 4.1. Parasitological findings

#### 4.1.1. Overall prevalence of Trypanosomosis

The current study revealed an overall trypanosomosis prevalence of 5.73% (22/384). Out of 384 animals examined 11 (2.87%) were infected with *T. vivax*, 9 (2.34%) were infected with *T. congolense* and 2 (0.52%) were mixed infected of both *T. vivax* and *T. congolense* (Table 1). This difference in trypanosomes species prevalence was not statistically significant (P>0.05).

**Table 1:** Overall prevalence of Trypanosoma species.

| Species | No. Positive | Prevalence (%) |
| --- | --- | --- |
| *T. vivax* | 11 | 2.87 |
| *T. congolense* | 9 | 2.34 |
| Mixed | 2 | 0.52 |
| **Total** | **22** | **5.73** |

#### 4.1.2 Prevalence of trypanosomosis with associated risk factors

In the present study, the association of prevalence with the various risk factors including kebele, age, sex, body condition and coat color were figured (Table 2). Trypanosomosis prevalence among kebeles was not statistically significant (P>0.05) with highest prevalence being recorded in Degaga Didesa followed by Oda Dideda and Meda Jalela respectively. Higher prevalence was observed in adult animals (>3 years old) than young (≤ 3 years) but the difference was not statistically significant. The prevalence of trypanosomosis was higher in females compared to males, but the difference was not statistically significant (P> 0.05). There was a significant variation in the prevalence of trypanosomosis (P<0.05) among cattle with different body conditions, poor body condition animals with the highest prevalence followed by medium and good body condition animals, respectively.

**Table 2:** Prevalence of trypanosomosis with associated risk factors.

| Risk factor | | Total examined | No. positive | Prevalence | $\chi^2$ (p- value) |
| --- | --- | --- | --- | --- | --- |
| Kebele | Meda Jalela | 132 | 6 | 4.5 % | 1.8995 (0.387) |
|  | Oda Dideda | 144 | 7 | 4.9 % |  |
|  | Degaga Didesa | 108 | 9 | 8.3 % |  |
| Age | $\leq 3$ years | 91 | 2 | 2.2% | 2.7537 (0.097) |
| | $>3$ years | 293 | 20 | 6.8% | |
| Sex | Male | 62 | 1 | 1.6% | 2.3196 (0.128) |
|  | Female | 322 | 21 | 6.5% |  |
| Body condition | Poor | 7 | 5 | 71% | 59.3481 (0.000) |
|  | Medium | 236 | 14 | 5.9% |  |
|  | Good | 141 | 3 | 2.1% |  |
| Coat color | Red | 299 | 20<br>1 1 0 0 | 6.7% | 2.8011 (0.592) |
|  | Black | 47 |  | 2.13% |  |
|  | Gray | 19 |  | 5.3% |  |
|  | Black spot | 9 |  | 0% |  |
|  | White | 10 |  | 0% |  |

The highest prevalence was recorded in cattle with red coat colour followed by gray and black coat colors, respectively. Trypanosoma was not recorded in black spot and white animals.

### 4.2. Haematological findings

The overall Packed Cell Volume (PCV) mean PCV value of 25.67 was recorded during the study period. The mean PCV value of infected animals was lower (20.18) than that of non-infected animals (26.0). The difference was statistically significant (P<0.05) (Table 3).

**Table 3:** Mean packed cell volume of infected and non-infected animals.

| Infection status | No. of animals | Mean ± SD | 95% CI |
| --- | --- | --- | --- |
| Parasitemic | 22 | 20.18 ± 2.59 | 19.03 - 21.33 |
| Aparasitemic | 362 | 26.01 ± 3.50 | 25.64 - 26.37 |
| **Total** | **384** | **25.67 ± 3.71** | **25.3 – 26.0** |

## 5. DISCUSSION

The results of the present study provide valuable insights into the prevalence of trypanosomosis among cattle, as well as its association with various risk factors and the impact on hematological parameters.

The overall prevalence of trypanosomosis was found to be 5.73%, indicating the presence of the disease in the study area. The present finding was comparable with the study conducted by [15] who reported an overall prevalence of 7.81 in their study on prevalence of bovine trypanasomosis in Guto Gida district of East Wollega Zone, Oromia Regional State, [16] who reported 8.96 % in their study on Epidemiology of bovine Trypanosomosis in Kamashi District of Benishangul Gumuz Regional State, Western Ethiopia, [17] whose finding showed 6% prevalence in his study on prevalence of bovine and host related risk factors in the neighboring Jawi district of the Amhara regional state south west of Ethiopia, [18] whose report showed 5.58 % prevalence in their study on trypanosomosis in Cattle Population of Pawi District of Benishangul Gumuz Regional State, Western Ethiopia, [19] who reported 5.43% prevalence in in their study on prevalence of bovine trypanosomosis and Apparent Density of Tsetse and Other Biting Flies in Dangur district of the Benishagul Gumuz region, western Ethiopia and [20] who reported 7.33% in their study on Prevalence of Bovine Trypanosomosis in Pawi District of the Benishngul Gumuz Region, North Western Ethiopia. However, the present finding was slightly lower than the studies made by [21] in the neighboring Dangur district who reported 11.27% in their study on trypanosomosis and its associated risks in cattle population, [18] who reported 13.30% in their study on Epidemiology of Cattle Trypanosomosis and Associated Anaemia in Mandura District of the Benishangul Gumuz regional state, west Ethiopia.

This prevalence is noteworthy as it highlights the extent of trypanosomosis among the examined cattle population. The distribution of trypanosome species showed that T. vivax, T. congolense, and mixed infections were identified, with T. vivax being the most prevalent species.

The prevalence rates for T. vivax, T. congolense, and mixed infections were provided, demonstrating a balanced distribution among the identified species. The lack of statistical significance in the differences between these species indicates that no particular trypanosome species predominates in the study area.

The study investigated the association of trypanosomosis prevalence with various risk factors. Kebele, age, sex, body condition, and coat color were considered as potential influencing factors. While kebele-wise prevalence did not show statistical significance, variations were observed concerning age, sex, body condition, and coat color. These findings suggest that certain factors such as age, body condition, and coat color may contribute to the susceptibility of cattle to trypanosomosis in the study area. This result agrees with previous reports [19-20]. This showed that animals with poor body condition are highly susceptible to trypanosomosis infection when compared with animals with good body condition.

The study included hematological assessments, focusing on the mean PCV values of infected and non-infected animals. Infected animals exhibited a significantly lower mean PCV (20.18) compared to non-infected animals (26.01). This is in line with previous results from different researchers [20, 22 - 24]. This finding is consistent with the expected impact of trypanosomosis on the hematological parameters of infected cattle. The lower PCV values in infected animals indicate anemia, a common consequence of trypanosome infections. Anemia, resulting from the destruction of red blood cells by the parasites, can lead to reduced productivity and overall well-being of the affected animals.

## 6. CONCLUSION AND RECOMMENDATIONS

In conclusion, this study provides valuable information on the prevalence of trypanosomosis, its association with various risk factors, and the hematological impact on infected cattle. The findings underscore the importance of targeted control measures and emphasize the need for further research to better understand the dynamics of trypanosomosis in the studied population.

## 7. ACKNOWLEDGEMENTS

We would like to acknowledge animal owners, district and Kebele officials for their corporations during sample collection.

## 8. AVAILABILITY OF DATA AND MATERIALS

The data sets used and/or analysed during the current study are available from the corresponding author on reasonable request.

## 9. ETHICS APPROVAL AND CONSENT TO PARTICIPATE

This research project was approved by the ethical review committee of the college of veterinary medicine and animal sciences, University of Gondar, Ethiopia. All efforts were made to minimize animal suffering during sample collection based on OIE standards. Informed oral consents were obtained from all animal owners who participated in the study.

## 10. CONSENT FOR PUBLICATION

Not applicable

## 13. AUTHORS’ CONTRIBUTIONS

Gemechis Yadesa: Data collection, Draft manuscript writing

Abebe Tesfaye Gessese: Proposal writing, Data analysis, manuscript writing and correction

